# Reconstitution of human cytochrome P450 activity using a *Leishmania* cell-free protein expression system

**DOI:** 10.1101/2025.06.20.660707

**Authors:** Wayne A. Johnston, Raine E. S. Thomson, Juan Alfaro-Palma, Robert E. Speight, Kirill Alexandrov, Elizabeth M. J. Gillam, James B. Y. Behrendorff

**Author notes:** Corresponding author: **Corresponding author James B. Y. Behrendorff** – Centre for Agriculture and the Bioeconomy, ARC Centre of Excellence in Synthetic Biology, School of Biology and Environmental Science, Queensland University of Technology, Brisbane, QLD 4000, Australia.

## Abstract

Cytochrome P450 enzymes (P450s) are ubiquitous in drug metabolism and natural product biosynthesis. Studying eukaryotic P450s has been limited by their dependence on membrane association and requirement for partner reductases. Here, we demonstrate cell-free synthesis and assay of human P450s 3A4 and 2D6 using *Leishmania tarentolae* translational extract. These P450s were co-expressed with various NADPH-cytochrome P450 reductases (CPRs), and activity was assayed directly using unpurified reactions. P450s 3A4 and 2D6 showed distinct preferences in reductase coupling: P450 3A4 activity was greatest when coupled to the human CPR, whereas P450 2D6 performed better when co-expressed with CPRs from *Arabidopsis thaliana*. Inhibition assays with chloramphenicol, terbinafine, and erythromycin yielded results consistent with known P450-drug interactions. We conclude that *Leishmania*-based cell-free protein synthesis resolves previous challenges in eukaryotic P450 expression, allowing for rapid and convenient functional studies of unmodified eukaryotic P450 systems, offering a practical tool for drug metabolism studies and biocatalyst discovery.

## Introduction

Cytochrome P450 enzymes (P450s) are heme monooxygenases that catalyse an extraordinary range of oxyfunctionalization reactions across all known domains of life^1^. P450s play major roles in xenobiotic metabolism (including metabolism of 80-90% of human and veterinary drugs and procarcinogens)^2^, and are required for biosynthesis of important secondary metabolites including sterols, alkaloids, carotenoids and more in animals, plants, fungi and microorganisms^3–5^. These activities make them attractive for biocatalysis. Therefore, robust methods for P450 expression and assay are sought both for research and industrial applications.

Cell-free protein synthesis (CFPS) is an attractive method for high-throughput expression, analysis and optimization of enzyme-catalyzed reactions prior to investment in whole-cell engineering and optimization of protein expression. CFPS systems can be prepared either from purified components (PURE systems) or from a fractionated cell-lysate (known as cell-free translational extract) that retains most of an organism’s soluble proteome and is supplemented with additional cofactors and enzymes to support transcription and translation^6,7^. Assay-ready enzymes can be synthesized in as little as two hours in microtiter formats amenable to automation^8,9^. Despite the potential utility of CFPS in P450 analysis and engineering, there are only a handful of reports where CFPS has been successfully applied to P450 production and analysis.

Expression of functional P450s requires supply of a heme cofactor and a partner reductase system to support the P450 catalytic cycle^10^. Bacterial P450s are generally soluble and are reduced by soluble redox mediators such as ferredoxins and flavodoxins, while most eukaryotic P450s and their cognate CPRs are anchored to the surface of the endoplasmic reticulum and typically require membrane association for correct folding and function^11^. Early studies of membrane anchoring domains from mammalian P450s and NADPH-cytochrome P450 reductases (CPRs) used wheat germ translational extracts for *in vitro* synthesis of proteins and analysis of membrane insertion^12,13^. These studies established *in vitro* functionality of membrane-targeting domains from a mammalian P450 and CPR but did not attempt to reconstitute enzymatic activity.

Cell-free P450 expression studies to-date have used a variety of approaches to supply the reducing power to the P450 enzyme. Soluble bacterial P450s TxtE^14^ (from *Streptomyces acidiscabies*) and BM3^15^ (from *Priestia megaterium*) were expressed using *E. coli*-derived translational extracts, while CypX from *Bacillus subtilis* was expressed in a translational extract derived from its native *B. subtilis* 164 host strain^16^. TxtE and CypX catalytic activities were supported by post-translational addition of a purified flavodoxin and flavodoxin reductase^14,16^. P450 BM3 naturally occurs as a fusion protein with its reductase domain and was functionally expressed from a single open reading frame^15^. The biosynthetic pathway for teleocidin B, including a C-N bond formation catalysed by P450 TleB from *Streptomyces eurocidicus*, was reconstituted in a translational extract derived from tobacco BY-2 cell suspension culture^17^. This CFPS system was chosen for its demonstrated capacity in producing functional non-ribosomal peptide synthases and high capacity for cofactor regeneration required for teleocidin B synthesis. In this example P450 activity was supported by addition of purified ferredoxin and ferredoxin reductase.

Active human P450s produced via CFPS were first demonstrated using a Chinese hamster ovary (CHO) cell line engineered to overexpress human CPR (hCPR), such that the overexpressed reductase was already present in microsomes upon preparation of the CHO cell-free translational extract^18^. More recently, functional co-expression of a plant P450 from *Arabidopsis thaliana* (cinnamate 4-hydroxylase) and its cognate reductase (Atr1) was reported using a wheat germ translational extract supplemented with synthetic liposomes, where reconstitution of enzyme activity was dependent on P450 and reductase co-expression for liposome co-insertion^19^. In *E. coli*, CHO, and wheat germ-based systems, P450 catalytic activity was enhanced by hemin supplementation, indicating limited availability of free heme in the translational extracts.

In this study we report co-expression of human P450s and reductases in a one-pot reaction using translational extract derived from *Leishmania tarentolae* (LTE). The quality of recombinant protein produced in LTE is comparable to other eukaryote-based CFPS that support full-length translation and low protein aggregation^20^. Unlike other eukaryotic hosts used for preparation of translational extracts,

*L. tarentolae* can be easily cultivated in standard shake flasks or bioreactors in low-cost microbial culture media^21^. LTE also contains residual membrane microsomes critical for functional expression of membrane-associated proteins^22^. Human P450s 3A4 and 2D6 were selected as models for this study due to their biomedical importance, together accounting for first pass metabolism of approximately 50% of drugs^2^. We demonstrate that full-length unmodified human P450s expressed in LTE are functional and that this format is suitable for combinatorial co-expression with alternative CPRs.

## Results and Discussion

### Establishment of P450 and CPR co-expression and functional assays

P450s and CPRs were encoded on separate plasmids and co-expressed in LTE. This co-expression strategy contrasts with previous cell-free P450 expression that relied on post-translational addition of purified reductases^14^ or engineered cell lines that overexpress a specific P450 reductase^18^. Human P450s 2D6 and 3A4 were expressed in LTE in combination with their cognate reductase (hCPR) or CPRs from *Arabidopsis thaliana* (Atr1 or Atr2). Additional hemin (for incorporation into P450s), flavin mononucleotide (FMN) and flavin adenine dinucleotide (FAD; both flavin cofactors being needed for incorporation into CPRs) were added post-translationally (see Methods). Analysis of BODIPY-lysine-labelled CFPS products on SDS-PAGE indicated two dominant product bands corresponding to the full-length P450s and CPRs, and few minor co-product bands (Figure 1A). A major fluorescent band corresponding to unincorporated BODIPY-lysine was observed in all lanes at approximately 15 kDa (Supporting Figure S1). Overall expression quality, determined as synthesis of full-length target proteins, was comparable to the best results achieved with LTE in a previous study benchmarking LTE against other CFPS systems^20^.

**Figure 1.**
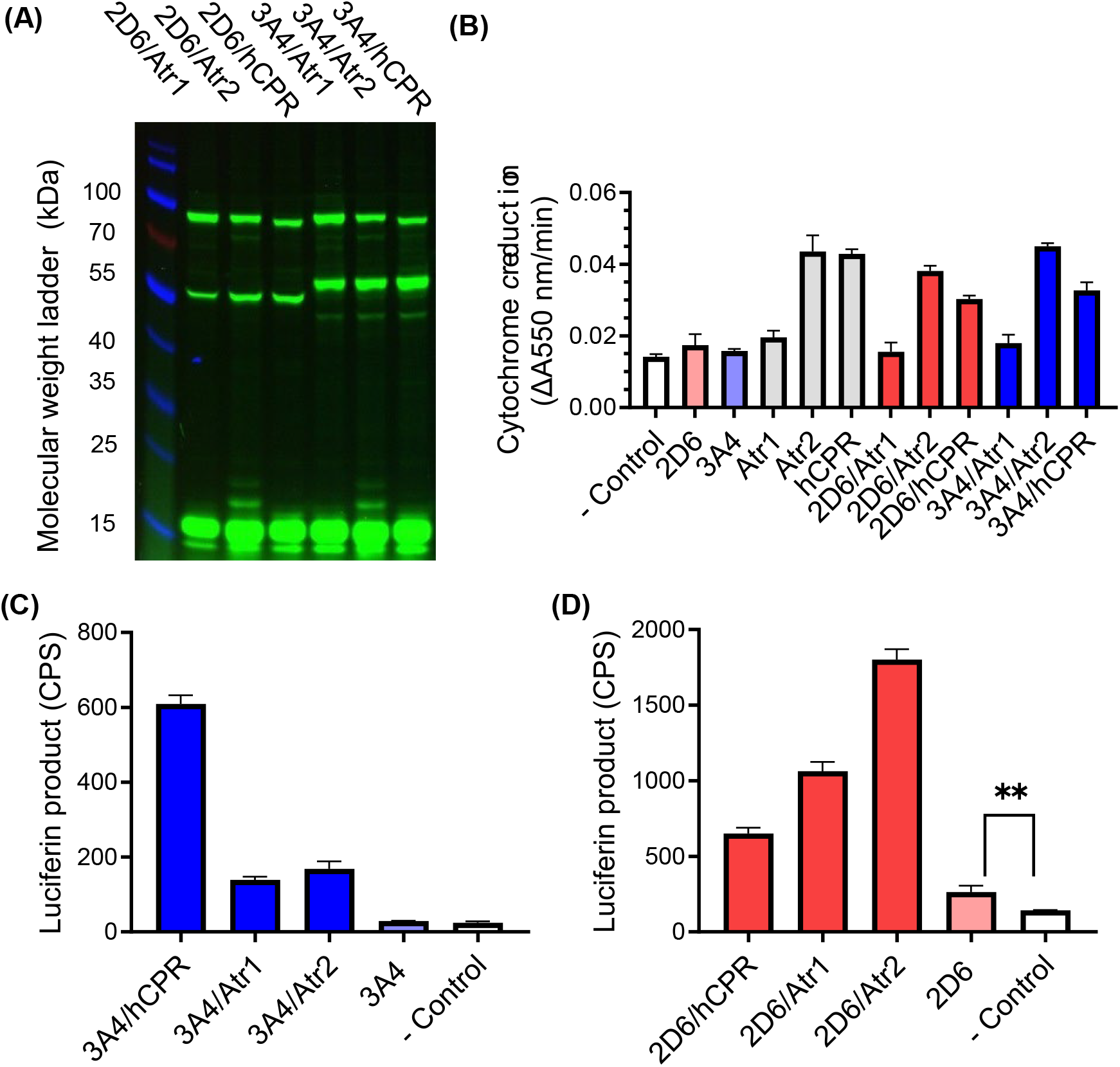
Co-expression and activity assays of cytochrome P450 enzymes and reductases in *Leishmania* translational extract. (A) Fluorescently labelled proteins synthesized in *Leishmania* translational extract were separated and visualised in SDS-PAGE. Anticipated molecular weights are: P450 2D6, 58 kDa; P450 3A4, 59 kDa; Atr1, 79 kDa; Atr2, 81 kDa; hCPR, 79 kDa. A protein molecular weight marker ladder is included in the left-hand lane. (B) Reduction of cytochrome *c* by CPR enzymes expressed independently (grey) or in co-expression with P450 2D6 (red) or P450 3A4 (blue) (n = 3 independent CFPS reactions, mean ± standard deviation). (C) Activity of P450 3A4 co-expressed with different reductases, incubated with the probe substrate Luciferin-IPA (n = 3 independent CFPS reactions, mean ± standard deviation). (D) Activity of P450 2D6 co-expressed with different reductases, incubated with probe substrate Luciferin-ME EGE (n = 3 independent CFPS reactions, mean ± standard deviation, ** = unpaired *t*-test p < 0.01).

Reductase activity was assayed using a colorimetric cytochrome *c* reduction assay where 2 µL of CFPS reaction product was diluted directly into assay buffer (Figure 1B). Human reductase hCPR and *A. thaliana* reductase Atr2 showed clear cytochrome *c* reduction activity when expressed individually or in combination with P450s. The rate of cytochrome *c* reduction by Atr1 was indistinguishable from the background rate of reduction in negative controls, though the published *k*_cat_ of cytochrome *c* reduction by Atr1 is less than 1% of that determined for Atr2 and hCPR^23^. Analysis of BODIPY-labelled Atr1 and Atr2 translation products indicated that Atr1 is translated at greater levels than Atr2, and that the lack of observable Atr1 activity in the cytochrome *c* assay is consistent with its known low catalytic turnover rather than poor expression (Supporting Figure S2).

Catalytic activity of P450s 2D6 and 3A4 was assayed using luminogenic probe substrates. P450s and CPRs were co-expressed by combining equimolar concentrations of P450 and CPR template DNA as this ratio was optimal for activity (Supporting Figure S3). Product formation by P450 3A4 was greatest when paired with its native reductase partner, hCPR, while P450 2D6 showed significantly greater activity with *A. thaliana* CPRs Atr2 and Atr1 (Figure 1C, D). Differences in coupling preference between P450s and CPRs have been reported previously^24,25^ and are likely due to structure-based differences in electron-transfer efficiency at the P450:CPR interface^26^. For example, product formation by *A. thaliana* P450 79A2 is faster when paired with Atr1^24^, despite the low catalytic rate of Atr1 relative to Atr2. In some cases P450s are non-functional when paired with the wrong reductase; PsiH from *Psilocybe cubensis* catalyses 4-hydroxylation of tryptamine when paired with its native CPR or with NCP1 from *Saccharomyces cerevisiae*, but has negligible activity when paired with Atr2^25^. Choice of CPR can also profoundly impact the product spectrum of catalytically promiscuous P450s^27^, presumably due to induced conformational changes in the P450 structure or differences in electron transfer rate.

A low level of P450 2D6 catalytic activity was observed even in the absence of CPR co-expression, suggesting that P450 2D6 may be reduced by low concentrations of endogenous *Leishmania* microsomal CPR^28^ or other reductants (Figure 1D, Supporting Figure S3). Marginal activity was observed when combining P450 and CPR from separate CFPS reactions but the greatest activity was observed when P450s and CPRs were co-expressed in a one-pot format (Supporting Figure S4), in agreement with reports of plant P450 and reductase co-expression in wheat germ translational extract^19^. Reductase-free catalysis, an emerging technology in P450 biocatalysis, was also demonstrated via CFPS of a P450 from a recently-reported ancestral reconstruction and validating functionality with its compatible reactive oxygen surrogate, bis(acetoxy)iodobenzene (Supporting Figures S5)^29^.

### Comparing functionality of P450s expressed in *L. tarentolae*- and *E. coli*-based CFPS

Heterologous expression of eukaryotic P450s in *E. coli* often requires modification of the N-terminal membrane anchor, but no universally successful rules have been derived to guide successful design despite an extensive history of investigation of P450 modifications to improve expression^24,30–33^. This study used full-length unmodified eukaryotic P450 and CPR coding sequences, which can be advantageous in the context of high-throughput protein expression as it eliminates resource-intensive design and testing of different N-terminal modifications for each enzyme. We compared P450 translation and functional assay between LTE and an *E. coli*-derived translational extracts.

No catalytic activity was observed when P450s 2D6 and 3A4 were co-expressed with hCPR in an *E. coli*-derived translational extract (Figure 2A). Translated proteins corresponding to P450 3A4 and hCPR were observed with *E. coli*-based CFPS, suggesting that the absence of catalytic activity may be due to protein misfolding or failure to reconstitute in a compatible membrane vesicle (Supporting Figure S1). No P450 2D6 translation product was observed with *E. coli*-based CFPS. This is consistent with previous studies of P450 2D6 expression in *E. coli* whole cells where modifications to the 2D6 coding sequence were required for protein translation^30^. Unexpectedly, combining P450 2D6 and hCPR template DNA in *E. coli* CFPS also prevented translation of hCPR, suggesting that the native P450 2D6 coding sequence inhibits protein translation. The reason for this inhibitory effect is unclear but could potentially be caused by stalled translation on the P450 2D6 mRNA, resulting in unproductive sequestration of *E. coli* ribosomes^34^. Despite the prevalence of PURE and *E. coli*-based CFPS technologies^35^, these are unlikely to be suitable for routine eukaryotic P450 screening.

**Figure 2.**
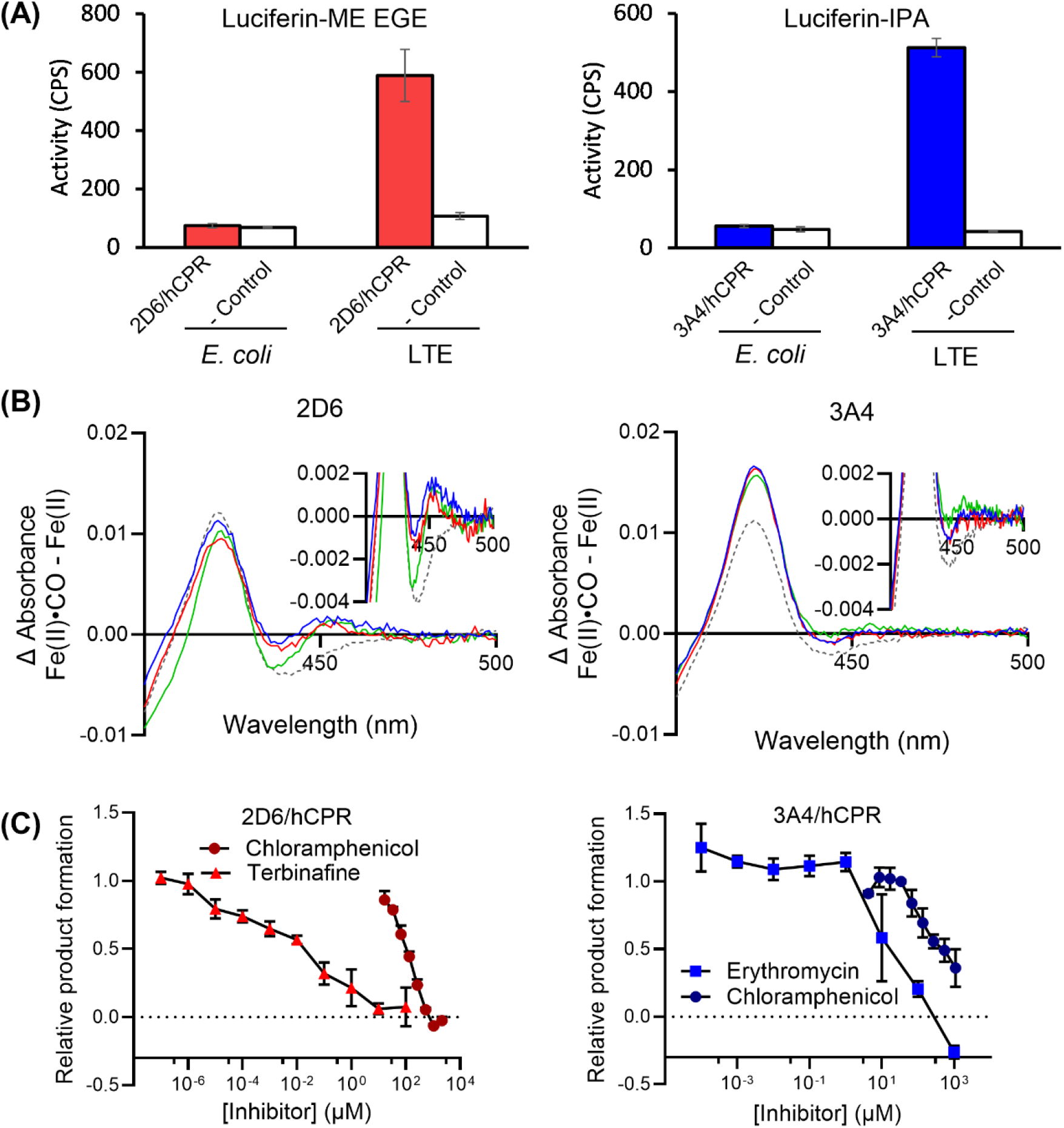
Comparison of P450 expression between LTE and *E. coli*-based CFPS, and evaluation of P450 performance in characteristic spectroscopic and enzyme inhibition assays. (A) P450s 2D6 or 3A4 were co-expressed with hCPR in LTE or *E. coli*-based CFPS. Luciferin product formation (luminescence counts per second) was recorded following incubations with Luciferin-ME EGE (for P450 2D6) or Luciferin-IPA (for P450 3A4) (n = 3 replicate CFPS reactions, mean ± standard deviation). (B) Fe(II)·CO *vs*. Fe(II) difference spectra of P450s 2D6 and 3A4 expressed in LTE (n = 3 independent diffusion-regenerated CFPS reactions, solid lines). Difference spectra of negative control reactions lacking template DNA but including hemin are shown with a grey dashed line (mean of n = 3 independent CFPS reactions). Inset plots show spectra expanded in the 450 nm region. (C) Inhibition of P450 2D6 by chloramphenicol (●, circles) or terbinafine (▴, triangles), and P450 3A4 by chloramphenicol (●, circles) or erythromycin (◼, squares). P450 activity is reported as luciferin product formation relative to luciferin formation in the absence of any inhibitor. All data represent n = 3 replicate assays (mean ± standard deviation).

### Characteristic ligand-binding spectra with carbon monoxide

Detection of the characteristic 450 nm absorbance peak by Fe(II).CO *vs*. Fe(II) difference spectroscopy is a reliable indicator of functional P450 expression^36,37^. Although standard 10 µL CFPS batch reactions were sufficient for activity screening, P450 expression titers were too low to detect the characteristic absorbance peak at 450 nm. Protein yield in CFPS can be enhanced by supplying additional nucleotides, amino acids, and energy cofactors via a dialysis membrane, increasing the yield and concentration of the protein product^38^. In this study we increased P450 expression yield via a simple diffusion-limited regeneration system where supplemental feed solution is added to a standard batch reaction without mixing, and slow diffusion of the feed solution occurs based on a density gradient (see Methods).

For each of P450 2D6 and 3A4, reaction products from ten CFPS reactions were pooled and the Fe(II) *vs*. Fe(II)·CO difference spectrum was recorded. A characteristic peak at 450 nm could be observed for P450 2D6 but not for 3A4 (Figure 2B). Although enzyme activity was readily detected for both enzymes, it is apparent that the P450 3A4 enzyme titer is below the detection limit of standard spectroscopic analysis. A large absorbance peak at 420 nm was attributed to excess unincorporated hemin (rather than ‘P420’, an inactive form of the enzyme), as this peak was also present in negative control CFPS reactions (free of template DNA) after hemin addition.

### Human P450 drug-inhibition assays

In addition to their central role in drug clearance, human P450s are also inhibited by many pharmaceuticals resulting in clinically significant drug-drug interactions^39^. P450 activity was assayed in the presence of known inhibitors of P450 2D6 (chloramphenicol^40^ and terbinafine^41^), and P450 3A4 (chloramphenicol^40^ and erythromycin^42^) (Figure 2C). Enzyme inhibition by chloramphenicol and terbinafine was typical of reversible competitive inhibitors. Erythromycin appeared to compete poorly against the substrate (3 µM Luciferin-IPA) until a threshold concentration of erythromycin was exceeded (> 1 µM) at which point P450 3A4 was strongly inhibited by further increases in erythromycin, consistent with the known competitive but irreversible mechanism of inhibition where erythromycin forms a stable intermediate metabolite complex with the reduced heme iron.

### Conclusion

Cell-free expression of eukaryotic P450s and their reductases was established with a leishmania-based translational extract. This protein expression format allowed for rapid and combinatorial co-expression of P450s and CPRs, and we demonstrated the utility of this system in identifying preferred P450-reductase pairings. No modifications to P450 protein sequences were needed for functional expression, and enzyme activity and inhibition assays required as little as 1 µL of CFPS reaction. This platform may be extended to enzyme screening for biocatalyst identification, optimization of reaction conditions and metabolic pathways. With widespread availability of low-cost synthetic DNA, CFPS may also enable rapid and affordable screening of P450 genetic variants for drug metabolism and drug-drug interactions, supporting safer and more personalized medicine.

## Materials and Methods

### Plasmid DNA

Genes encoding P450s and CPRs were synthesised by GeneUniversal (http://www.geneuniversal.com) and cloned into the pCellFree plasmid vector containing the species-independent translation initiation sequence (SITS)^43^ for use in LTE and other cell-free protein expression systems. Further sequence details are included in the Supporting Information (Appendix S1).

### Batch cell-free protein synthesis

LTE cell-free lysate was produced from *L. tarentolae* using previously developed protocols^44^. All assays except CO-difference spectrum measurement used a standard batch reaction format (10 µL, 2 h incubation 25 °C) in PCR tubes or 384-well microtiter plates. Protein synthesis products were either used immediately for P450 activity assays or snap frozen in liquid nitrogen and stored at −80 °C for later use. DNA template input was set to 25 nM of purified plasmid. Co-expression of P450s and CPRs used 12.5 nM P450 and 12.5 nM CPR template DNA unless stated otherwise. Negative control reactions lacked template DNA.

For visualisation of protein expression on SDS-PAGE (4-20% acrylamide, Invitrogen NW04125), fluorescent BODIPY-lysine tRNA (FluoroTect, Promega, L5001) was added to protein synthesis reactions (1:100, v/v). Translation products were analysed alongside a molecular weight marker ladder (ThermoScientific PageRuler Prestained Protein Ladder, ThermoScientific Cat #26616).

### Diffusion regenerated cell-free protein synthesis

For spectral assays of P450s, CFPS yield was increased via diffusion of additional regeneration solution into the batch synthesis reaction. Batch reactions were setup as above, and then 5 µL was carefully pipetted (without mixing) to the base of a 384 well plate well (Optiplate, ThermoFisher, 6007290) already containing 45 µL of regeneration solution. The regeneration solution contained small molecule substrates equimolar to those in the cell-free batch reaction (1.7 mM ATP, 0.5 mM GTP, 0.5 mM CTP, 0.5 mM UTP, 5 mM magnesium acetate, 0.25 mM spermidine, 2 mM dithiothreitol, 40 mM creatine phosphate, 20 mM HEPES-KOH pH 7.6, polyethylene glycol 3000 (2%, v/v), 1× protease inhibitor (cOmplete EDTA-free, Roche, 11873580001), 0.14 mM of each amino acid). Diffusion-regenerated reactions were incubated for 5 h at 25 °C without mixing and then harvested by carefully pipetting the reaction back out of the plate well base (i.e. with minimal mixing with the regeneration solution). A volume equivalent to 1.5-fold the volume (7.5 µL) of completed reaction was harvested to help compensate for inevitable mixing of reaction and regeneration solutions during incubation.

### Fe(II).CO *vs*. Fe(II)-difference spectroscopy

For Fe(II).CO *vs*. Fe (II)-difference spectroscopy, ten 7.5 µL harvested diffusion-regenerated expressions of P450 2D6 or 3A4 as above were combined (75 µL *in toto*) in a single clear 384-well plate well for each triplicate spectral measurement, with Fe(II).CO *vs*. Fe (II)-difference spectra measured similarly to those previously described for whole cell P450 measurements^45^. Hemin (Sigma-Aldrich, 51280) was added to CFPS products (2 µM, incubated for 20 minutes at room temperature) and samples were reduced via addition of sodium dithionite (10 mg/mL incubated for 10 minutes at room temperature). Absorbance spectra of reduced samples were recorded (400-500 nm at 1 nm intervals) (Varioskan LUX multimode plate reader, Thermo Fisher). Samples were then subjected to a 100% CO atmosphere for 10 minutes in a dedicated gassing box, and absorbance spectra of reduced CO-treated samples were recorded. Difference spectra were calculated by subtracting the initial reduced sample absorbance spectra from the CO-treated sample spectra.

### Reductase and cytochrome P450 activity assays

Reductase activity was measured as cytochrome *c* reduction^36^ in a clear 384-well plate. The completed batch expression reaction (2 µL) was incubated for 5 minutes at room temperature with 32 µL 50 µM cytochrome *c* and 2 µL cofactor mix (10 µM of both FAD and FMN). After preheating the plate at 37 °C, reactions were started via addition of 4 µL NADPH regeneration solution (NGS) comprising 100 mM glucose-6-phosphate, 2.5 mM NADP^+^ and 10 U/ml glucose-6-phosphate dehydrogenase. Reductase activity was measured as the maximum rate of increase in absorbance at 550 nm.

For measurement of P450 activity, luciferin-based substrate kits were used that were specific for P450 2D6 (Luciferin-ME EGE, Promega, P450Glo V8891) and P450 3A4 (Luciferin-IPA, Promega, P450Glo, V9001) and reactions were carried out in 384-well microtitre plates (Optiplate, Thermo Fisher, 6007290). Batch expression reaction (1 µL) was added to 1 µL cofactor mix (8 µM FAD, 8 µM FMN, 20 µM hemin) and 6 µL phosphate-buffered saline. Reactions were initiated with addition of 1 µL NGS plus 1 µL luciferin substrate (to the concentrations suggested by the manufacturer: 3 µM Luciferin-IPA; 30 µM Luciferin-ME EGE). Reactions were incubated at 37 °C for 1 h prior to addition of luciferin detection reagent according to the manufacturer’s instructions. Luminescence was measured in a multimode microplate reader (Spark, Tecan) for 20 minutes. P450 activity was reported as the maximum luminescence signal in counts per second (CPS).

For preparation of P450 inhibition curves, the luminescent P450-Glo assays were used as above, with the addition of known inhibitors (chloramphenicol, P450s 2D6 and 3A4; terbinafine, P450 2D6; erythromycin, P450 3A4). Inhibitors were added to incubations with P450s as serial dilutions with vehicle as diluent (chloramphenicol, 5% DMSO; terbinafine and erythromycin, 5% ethanol). P450 activity was calculated by subtracting the background signal observed in negative control incubations (where template DNA had been omitted) and normalising to signal observed in vehicle controls (5% v/v DMSO or 5% v/v ethanol).

## Author information

### Author contributions

W.A.J., R.E.S.T, J. A.-P. and J. B. Y. B. contributed the experimental work. W. A. J., E. M. J. G., K. A., and J. B. Y. B. conceived of and designed the study. Project funding was secured by R. E. S. All authors contributed to discussion of experimental results and preparation of the manuscript.

### Notes

The authors declare no competing financial interest.

### Supporting information

Supporting information file 1: Protein sequence data, additional experimental methods, and additional results related to optimisation of P450 CFPS expression and assay.

## Acknowledgements

This work was supported by the Australian Research Council Centre of Excellence in Synthetic Biology (project CE200100029).

