## Supporting Information for "Reconstitution of human cytochrome P450 activity using a *Leishmania* cell-free protein expression system"

**Appendix S1 Peptide sequences encoded in pCellFree expression plasmid**

All protein coding sequences cloned in the pCellFree vector are preceded by a short sequence of mRNA hairpins that form part of the species independent translation sequence (SITS) described by Mureev *et al*. (2009). This leader sequence is translated in-frame with the protein of interest, resulting in the following N-terminal extension: MTVMYKVCKDIKHVSET.

>P450_2D6

MTVMYKVCKDIKHVSETMGLEALVPLAVIVAIFLLLVDLMHRRQRWAARYPPGPLPLPGLGNLL
HVDFQNTPYCFDQLRRRFGDVFSLQLAWTPVVVLNGLAAVREALVTHGEDTADRPPVPITQILG
FGPRSQGVFLARYGPAWREQRRFSVSTLRNLGLGKKSLEQWVTEEAACLCAAFANHSGRPFR
PNGLLDKAVSNVIASLTCGRRFEYDDPRFLRLLDLAQEGLKEESGFLREVLNAVPVLLHIPALAG
KVLRFQKAFLTQLDELLTEHRMTWDPAQPPRDLTEAFLAEMEKAKGNPESSFNDENLRIVVADL
FSAGMVTTSTTLAWGLLLMILHPDVQRRVQQEIDDVIGQVRRPEMGDQAHMPYTTAVIHEVQR
FGDIVPLGVTHMTSRDIEVQGFRIPKGTTLITNLSSVLKDEAVWEKPFRFHPEHFLDAQGHFVKP
EAFLPFSAGRRACLGEPLARMELFLFFTSLLQHFSFSVPTGQPRPSHHGVFAFLVSPSPYELCAV
PR

>P450_3A4

MTVMYKVCKDIKHVSETMALIPDLAMETWLLLAVSLVLLYLYGTHSHGLFKKLGIPGPTPLPFLG
NILSYHKGFCMFDMECHKKYGKVWGFYDGQQPVLAITDPDMIKTVLVKECYSVFTNRRPFGPV
GFMKSAISIAEDEEWKRLRSLLSPTFTSGKLKEMVPIIAQYGDVLVRNLRREAETGKPVTLKDVFG
AYSMDVITSTSFGVNIDSLNNPQDPFVENTKKLLRFDFLDPFFLSITVFPFLIPILEVLNICVFPREV
TNFLRKSVKRMKESRLEDTQKHRVDFLQLMIDSQNSKETESHKALSDLELVAQSIIFIFAGYETTS
SVLSFIMYELATHPDVQQKLQEEIDAVLPNKAPPTYDTVLQMEYLDMVVNETLRLFPIAMRLERV
CKKDVEINGMFIPKGVVVMIPSYALHRDPKYWTEPEKFLPERFSKKNKDNIDPYIYTPFGSGPRN
CIGMRFALMNMKLALIRVLQNFSFKPCKETQIPLKLSLGGLLQPEKPVVLKVESRDGTVSGA

>hCPR

MTVMYKVCKDIKHVSETMADSHVDTSSTVSEAVAEEVSLFSMTDMILFSLIVGLLTYWFLFRKKK
EEVPEFTKIQTLTSSVRESSFVEKMKKTGRNIIVFYGSQTGTAEEFANRLSKDAHRYGMRGMSAD
PEEYDLADLSSLPEIDNALVVFCMATYGEGDPTDNAQDFYDWLQETDVDLSGVKFAVFGLGNK
TYEHFNAMGKYVDKRLEQLGAQRIFELGLGDDDGNLEEDFITWREQFWPAVCEHFGVEATGE
ESSIRQYELVVHTDIDAAKVYMGEMGRLKSYENQKPPFDAKNPFLAAVTTNRKLNQGTERHLM
HLELDISDSKIRYESGDHVAVYPANDSALVNQLGKILGADLDVVMSLNNLDEESNKKHPFPCPT
SYRTALTYYLDITNPPRTNVLYELAQYASEPSEQELLRKMASSSGEGKELYLSWVVEARRHILAIL
QDCPSLRPPIDHLCELLPRLQARYYSIASSSKVHPNSVHICAVVVEYETKAGRINKGVATNWLRA
KEPAGENGGRALVPMFVRKSQFRLPFKATTPVIMVGPGTGVAPFIGFIQERAWLRQQGKEVGET
LLYYGCRRSDEDYLYREDVAQFHRDGALTQLNVAFSREQSHKVYVQHLLKQDREHLWKLIEGG
AHIYVCGDARNMARDVQNTFYDIVAELGAMEHAQAVDYIKKLMTKGRYSLDVWS

>Atr1

MTVMYKVCKDIKHVSETMTSALYASDLFKQLKSIMGTDSLSDDVVLVIATTSLALVAGFVVLLWKK
TTADRSGELKPLMIPKSLMAKDEDDDLDLGSGKTRVSIFFGTQTGTAEGFAKALSEEIKARYEKAA
VKVIDLDDYAADDDQYEEKLKKETLAFFCVATYGDGEPTDNAARFYKWFTEENERDIKLQQLAY
GVFALGNRQYEHFNKIGIVLDEELCKKGAKRLIEVGLGDDDQSIEDDFNAWKESLWSELDKLLK
DEDDKSVATPYTAVIPEYRVVTHDPRFTTQKSMESNVANGNTTIDIHHPCRVDVAVQKELHTHE
SDRSCIHLEFDISRTGITYETGDHVGVYAENHVEIVEEAGKLLGHSLDLVFSIHADKEDGSPLESA
VPPPFPGPCTLGTGLARYADLLNPPRKSALVALAAYATEPSEAEKLKHLTSPDGKDEYSQWIVAS
QRSLLEVMAAFPSAKPPLGVFFAAIAPRLQPRYYSISSSPRLAPSRVHVTSALVYGPTPTGRIHKG
VCSTWMKNAVPAEKSHECSGAPIFIRASNFKLPSNPSTPIVMVGPGTGLAPFRGFLQERMALKE
DGEELGSSLLFFGCRNRQMDFIYEDELNNFVDQGVISELIMAFSREGAQKEYVQHKMMEKAA
QVWDLIKEEGYLYVCGDAKGMARDVHRTLHTIVQEQEGVSSSEAEAIVKKLQTEGRYLRDVW

>Atr2

MTVMYKVCKDIKHVSETMSSSSSSSTSMIDLMAAIIKGEPVIVSDPANASAYESVAAELSSMLIEN
RQFAMIVTTSIAVLIGCIVMLVWRRSGSGNSKRVEPLKPLVIKPREEEIDDGRKKVTIFFGTQTGTA
EGFAKALGEEAKARYEKTRFKIVDLDDYAADDDEYEEKLKKEDVAFFFLATYGDGEPTDNAARFY
KWFTEGNDRGEWLKNLKYGVFGLGNRQYEHFNKVAKVVDDILVEQGAQRLVQVGLGDDDQC
IEDDFTAWREALWPELDTILREEGDTAVATPYTAAVLEYRVSIHDSEDAKFNDINMANGNGYTVF
DAQHPYKANVAVKRELHTPESDRSCIHLEFDIAGSGLTYETGDHVGVLCDNLSETVDEALRLLD
MSPDTYFSLHAEKEDGTPISSSLPPPFPPCNLRTALTRYACLLSSPKKSALVALAAHASDPTEAER
LKHLASPAGKDEYSKWVVESQRSLLEVMAEFPSAKPPLGVFFAGVAPRLQPRFYSISSSPKIAET
RIHVTCALVYEKMPTGRIHKGVCSTWMKNAVPYEKSENCSSAPIFVRQSNFKLPSDSKVPIIMIG
PGTGLAPFRGFLQERLALVESGVELGPSVLFFGCRNRRMDFIYEEELQRFVESGALAELSVAFSR
EGPTKEYVQHKMMDKASDIWNMISQGAYLYVCGDAKGMARDVHRSLHTIAQEQGSMDSTKA
EGFVKNLQTSGRYLRDVW

>2CE

MTVMYKVCKDIKHVSETMAKKTSSKGKLPPGPTPLPIIGNILQLNTKNLPKSLHKLSEKYGPVFTL
YLGSQRVVVLHGYEAVKEALIDHGDEFSGRGNMPIIDKINKGLGIVFSNGERWKQLRRFALTTL
RNFGMGKKSIEERIQEEAQYLVEELRNTKGQPFDPTFLLSCAVSNVICSIVFGKRFDYEDKKFLTL
MNLLNENFRLLNSPWGQLYNLFPSLMDYLPGPHHKIFKNFEELKDFILERVKEHQETLDPNSP
RDFIDCFLIKMEQEKQNPKSEFNMENLVMTTLDLFSAGTETTSTTLRYGLLILLKYPEIEEKVHEEI
DRVIGRNRSPCMADRSQMPYTDAVIHEIQRFIDLVPLGLPHAVTQDTHFRQYIIPKGTTIFPLLSS
VLHDSKEFPNPEQFNPGHFLDENGSFKKSDYFMPFSAGKRICVGEGLARMELFLFLTTILQNFT
LKPLVDPKDIDITPEVSGFGNVPRPYQLCFLPRSTHHHHHH

### Appendix S2 Supporting methods

*P450 activity assay supported by reactive oxygen surrogate*

P450 2CE[1,2] or EGFP was expressed in standard 10 µL CFPS batch reactions in LTE. CFPS reactions were diluted to 100 µL in 50 mM potassium phosphate buffer (pH 7.4), 10 µM hemin, and 100 µM Luciferin multi-CYP substrate (Promega). Reactions were initiated by adding BAIB to a final concentration of 1 mM. Reactions were incubated for 1 h at 37 °C prior to measuring luciferin formation according to the methods provided in the Promega P450Glo assay manual.

Supporting Figure S1

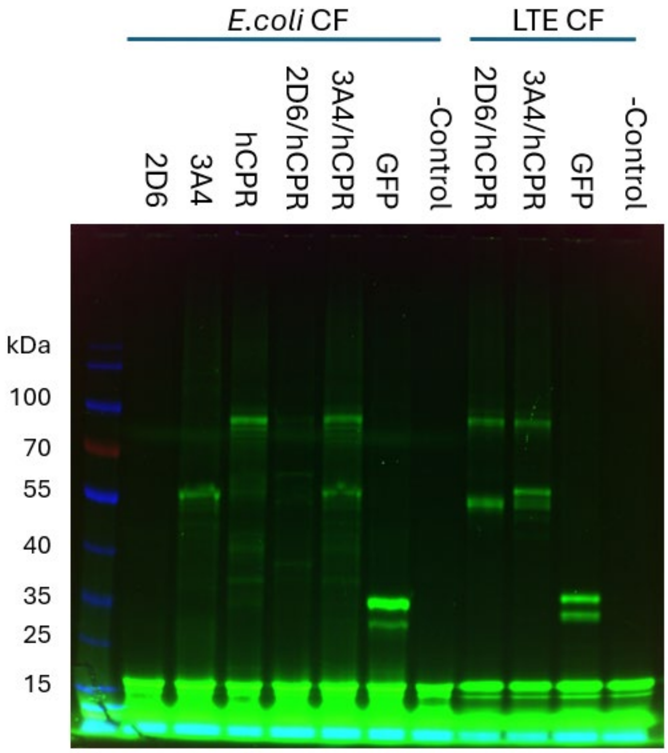


**Supporting Figure S1 Translation products from CFPS of P450s and hCPR using *E. coli* and *L. tarentolae* translational extracts.** Fluorescent BODIPY-lysine labelled proteins synthesised in *E. coli* and *L. tarentolae* translational extracts were separated and visualised in SDS-PAGE. Template DNA encoding P450 2D6, P450 3A4, hCPR, or EGFP was added to *E. coli* or *L. tarentolae* translational extracts and incubated for 2 h at 25 °C. Template DNA for P450s and hCPR were combined for co-expression (indicated by 2D6/hCPR and 3A4/hCPR). Negative controls (-Control) indicate incubations of translational extract with BODIPY-lysine but without addition of template DNA. A protein molecular weight marker ladder is included in the left-hand lane.

Supporting Figure S2

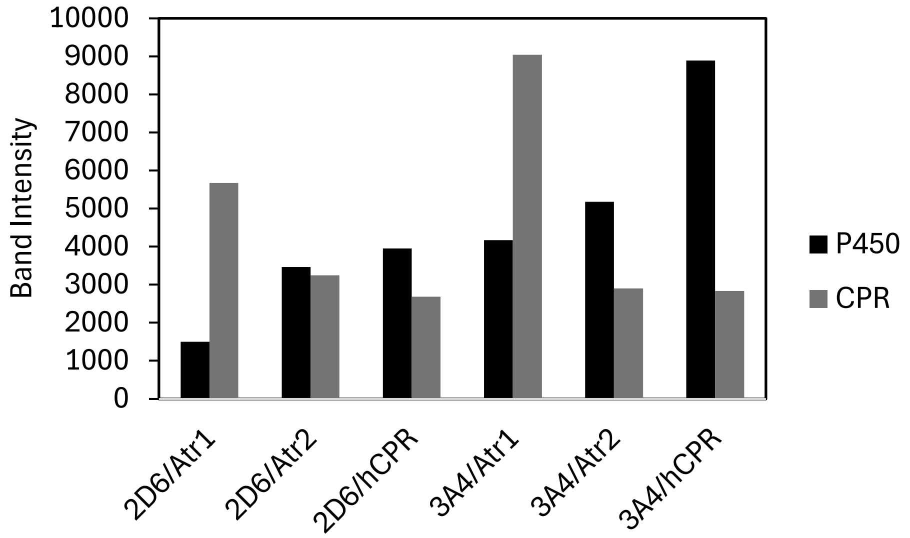


**Supporting Figure S2 Relative yield of target proteins from cell-free protein synthesis.** Fluorescence intensity of protein products resolved on SDS-PAGE in Figure 1A was quantified (ImageLab software, Bio Rad Inc.).

Supporting Figure S3

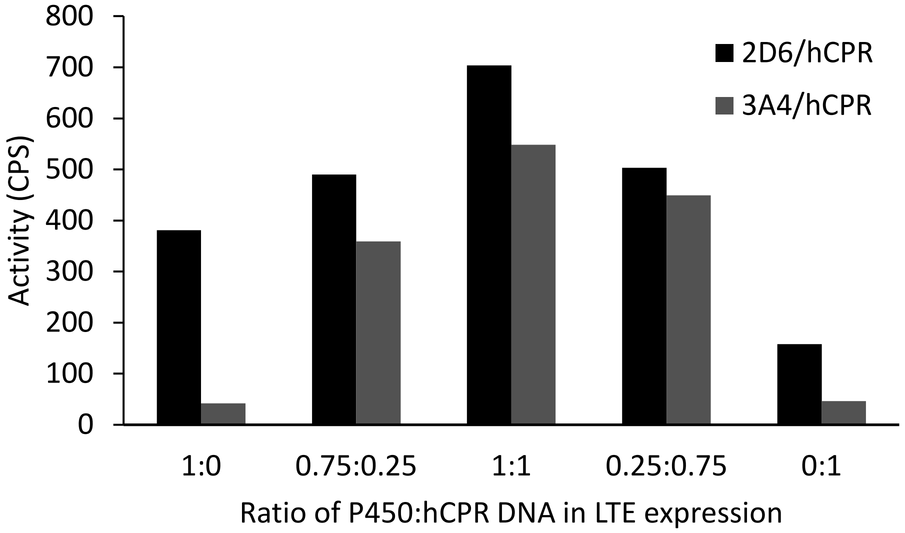


**Supporting Figure S3 P450 and CPR co-expression with different ratios of template DNA.** P450s 2D6 and 3A4 were co-expressed with hCPR by adding different ratios of template DNA in Leishmania translational extract. P450 2D6 activity was assayed with Luciferin-ME EGE; P450 3A4 activity was assayed with Luciferin-IPA.

Supporting Figure S4

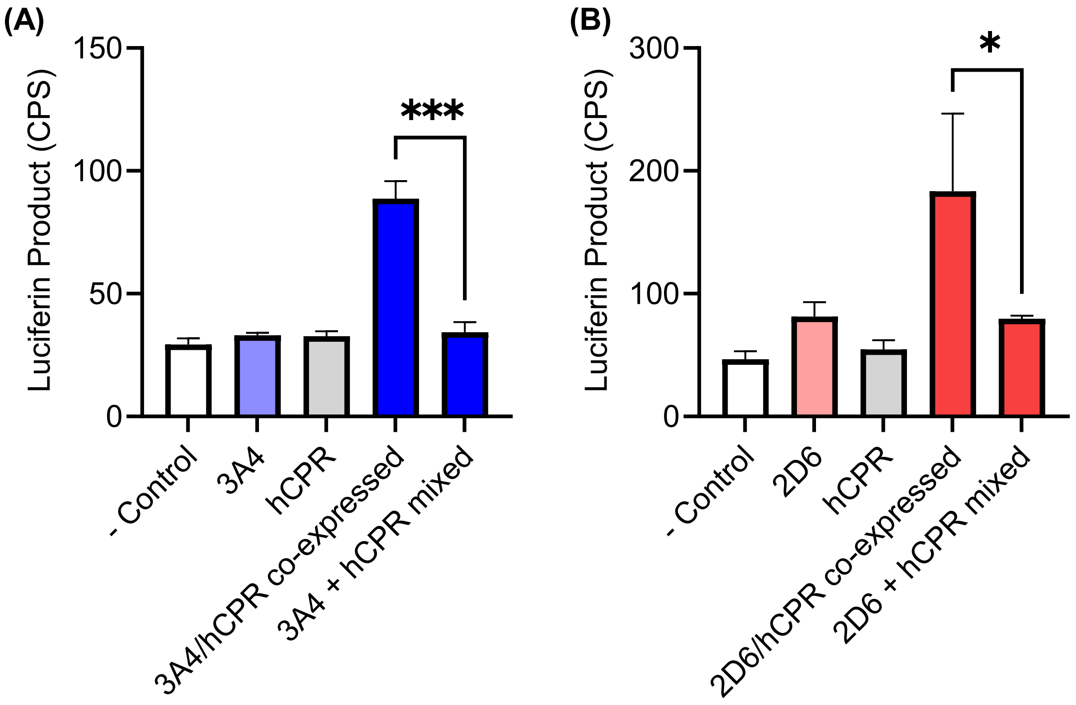


**Supporting Figure S4 Activity of P450s co-expressed with hCPR or combined post-translationally.** P450 2D6 and P450 3A4 were each co-expressed with human reductase (hCPR) in a single CFPS reaction, or expressed individually and then combined post-translation. P450 activity was assayed using the substrate Luciferin-IPA (P450 3A4) or Luciferin-ME EGE (P450 2D6). Data represent the mean + standard deviation of n = 3 independent CFPS reactions and subsequent enzyme assays. Asterisks indicate statistically significant differences at the $p < 0.05$ (*) and $p < 0.005$ (***) level, using an unpaired Student's *t*-test.

Supporting Figure S5

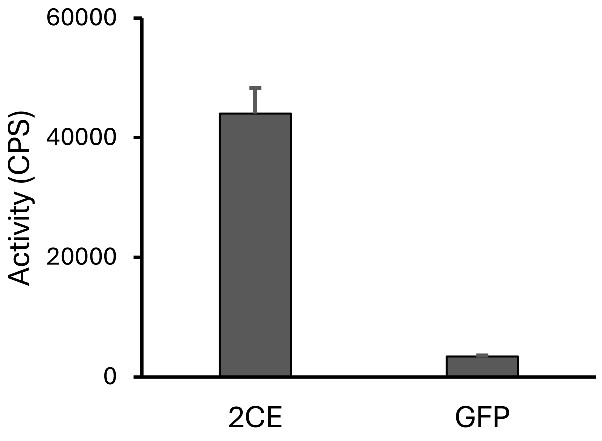


**Supporting Figure S5 P450 activity supported by reactive oxygen surrogate bi(trifluoroacetoxy)iodobenzene.** Synthetic ancestral P450 2CE[1,2] was expressed in Leishmania translational extract and incubated with the luminogenic substrate, Luciferin multi-CYP, and 1 mM bi(trifluoroacetoxy)iodobenzene. GFP was expressed and assayed as a negative control. Data represent the mean + standard deviation from n = 3 independent CFPS reactions and subsequent enzyme assays.

(1) Thomson, R. E. S. Structural and Functional Characterisation of Ancestral Cytochromes P450 from Family 2 in Tetrapods., The University of Queensland, 2021. https://doi.org/10.14264/a159633 (accessed 2025-06-13).
(2) Strohmaier, S. J.; Baek, J. M.; De Voss, J. J.; Jurva, U.; Andersson, S.; Gillam, E. M. J. An Inexpensive, Efficient Alternative to NADPH to Support Catalysis by Thermostable Cytochrome P450 Enzymes. *ChemCatChem* **2020**, *12* (6), 1750–1761. https://doi.org/10.1002/cctc.201902235.
